# Environmental DNA (eDNA) sampling strategies influence estimates of freshwater fish eDNA concentrations

**DOI:** 10.1101/2025.09.05.674411

**Authors:** Vautier Marine, Bylemans Jonas, Baudoin Jean-Marc, Guillard Jean, Goulon Chloé, Logez Maxime, Isabelle Domaizon

**Author notes:** Shared first author (Vautier Marine, Bylemans Jonas).

## Abstract

Quantifying *in-situ* environmental DNA (eDNA) concentrations is increasingly used to infer fish abundance and biomass in freshwater ecosystems. While various sampling strategies (i.e. the combination of sample collection, eDNA preservation and extraction protocols) have been proposed to collect eDNA, little efforts have been undertaken to evaluate how these different strategies affect taxon-specific eDNA recovery and the subsequent estimates of *in-situ* eDNA concentrations. In this study, we compared a point (i.e. eDNA collection from relatively small and spatially separated water samples filtered using low capacity filter units) and a spatially integrated sampling strategy (i.e. eDNA collected from spatially integrated larger water volumes filtered using high capacity filter units) in two natural lakes. Through quantitative analyses of total DNA, total fish eDNA and species-specific eDNA, we assessed the performance of both strategies to infer *in-situ* eDNA concentrations. Our results showed that the integrated strategy led to a reduced recovery of fish eDNA and a subsequent underestimation of *in-situ* eDNA concentrations compared to the point sampling strategy. While the exact mechanisms underlying this pattern require further investigation, our findings highlight the importance of carefully selecting sampling strategies according to study objectives.

## INTRODUCTION

Environmental DNA (eDNA) analyses have revolutionized biodiversity monitoring in aquatic ecosystems (Deiner et al., 2017). A critical first step in all eDNA surveys is to collect and capture eDNA from the environmental matrix. Several comparative studies have assessed the impact of individual steps in the overall sampling strategy (i.e. eDNA sampling, preservation or extraction) and have reported effects on DNA recovery and detection probabilities (Hinlo et al., 2017; Spens et al., 2016). Different filter types have also shown to affect the correlation between *in-situ* eDNA concentrations and species abundances or biomass (Eichmiller et al., 2016; Lacoursière-Roussel et al., 2016; Takahashi et al., 2020). While disentangle the effects of the individual components of the sampling workflow allows to optimize protocols; from a practical perspective, alternative sampling strategies often rely on a combination of eDNA sampling, preservation and extraction protocols.

Within the current eDNA literature, surprisingly few studies have directly compared alternative sampling strategies in terms of eDNA recovery and estimates of *in-situ* eDNA concentrations. Most current studies aiming to quantify *in-situ* eDNA concentrations in freshwater ecosystems have relied on collecting spatially separated point samples (i.e. point sampling (PS) strategy), which involves filtering small water volumes (*ca*. 1-2 L) using low capacity filter capsules (e.g. Lacoursière-Roussel et al. (2015); Takahara et al. (2012) and Yates et al. (2023)). An alternative sampling strategy has also been developed where larger volumes of water (*ca*. 30-45 L) are filtered using high-capacity filters along spatially integrated transects (i.e. integrated sampling (IS) strategy). While these integrated strategies were initially designed for qualitative community surveys in lakes and rivers (Civade et al., 2016; Hervé et al., 2022), several recent studies have used them for quantitative eDNA surveys (Condachou et al., 2024; Hervé et al., 2023; Pont et al., 2018). Comparisons between the two above-mentioned sampling strategies are limited and existing studies have focused solely on the difference in sampling volumes and neglected the ability to obtain spatially integrated information when using the IS strategy (Peixoto et al., 2021, 2023). This earlier work has shown that higher water volumes, filtered with high-capacity filters, collected more total DNA but yielded lower species-specific detection probabilities and lower eDNA concentrations compared to smaller water volumes collected using low capacity filters (Peixoto et al., 2021, 2023). Consequently, an integrated strategy may enhance total DNA recovery but may have a reduced recovery of taxon-specific eDNA estimates.

The current lack of comparative studies between the point and integrated sampling strategies hinders the ability to make informed decisions when designing eDNA sampling campaigns in freshwater ecosystems. To address this knowledge gap, we implemented both strategies in two natural lakes and we evaluated the effectiveness of both strategies in terms of eDNA recovery (i.e. total DNA and fish eDNA standardized by total DNA or total water volume filtered) and the quantitative estimates of *in-situ* eDNA concentrations for five fish species.

## MATERIAL AND METHODS

### Sampling

We selected two natural lakes in France as sampling sites, differing in size and trophic state (see Figure 1). Lake Bourget, one of the largest natural lake in France (*ca*. 4,397 ha.), is oligotrophic and reaches a maximum depth of *ca*. 145 m (Jenny et al., 2024). In contrast, Lake Paladru is smaller (*ca*. 355 ha.), shallower (maximum depth is *ca*. 36 m), and classified as mesotrophic (STE, 2024).

**Figure 1.**
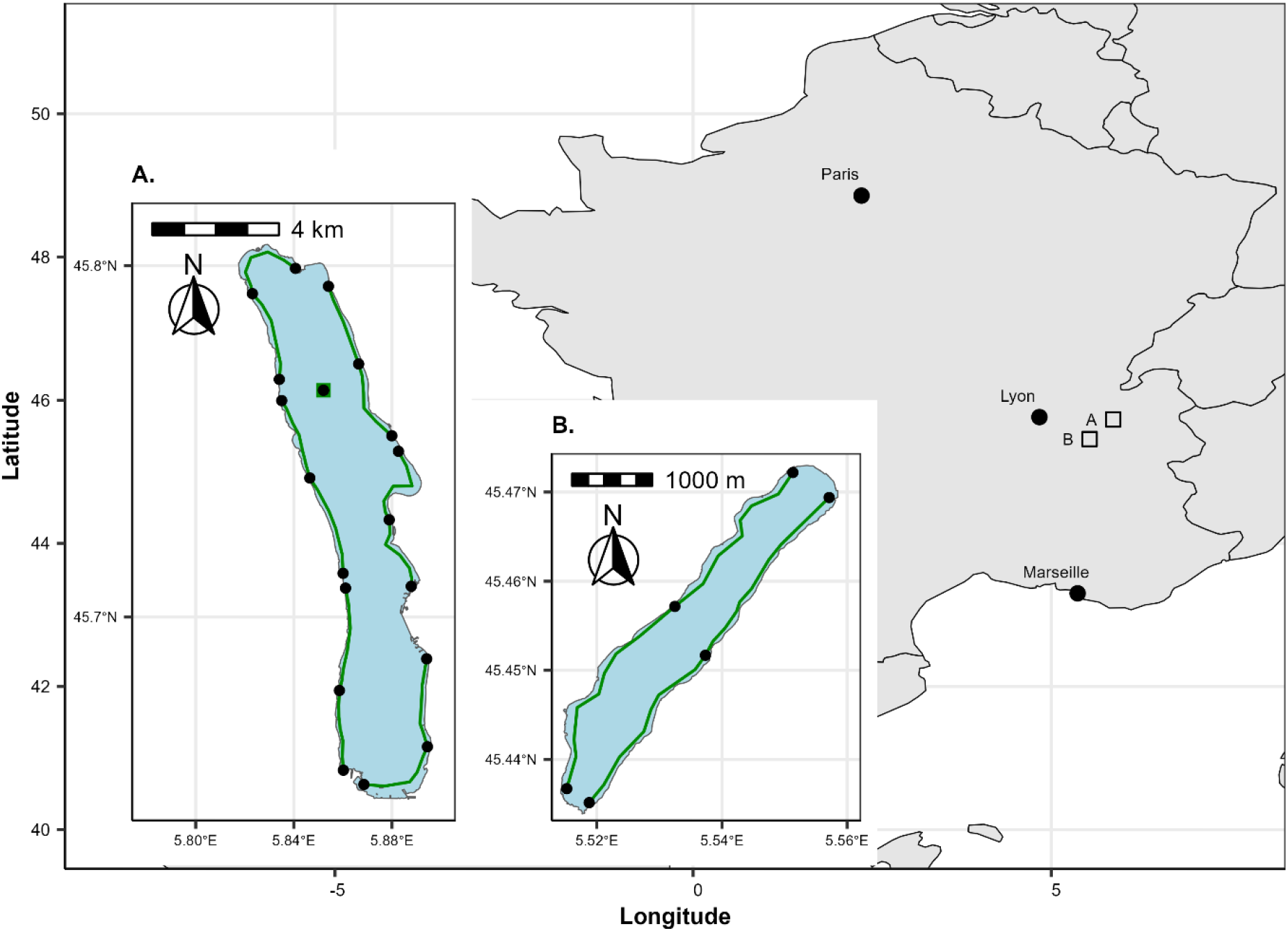
Geographic location of the two lakes and a visual representation of the environmental DNA sampling strategies for Lake Bourget (A) and Paladru (B). Sampling consisted of spatially integrated sub-surface samples collected using high capacity Waterra filters (green lines: 6 transects for Lake Bourget, 2 transects for Lake Paladru), a deep-water sample collected using a Waterra filter in Lake Bourget (green square), and sub-surface point samples collected using Sterivex filters (black dots: 3 samples per transect and deep-water sample).

All material for the collection and processing of eDNA samples were single-use or thoroughly decontaminated using an acid-based washing cycle with a Lancer Ultima Series Laboratory Washer followed by a hydrogen peroxide treatment (Vautier & Galiegue, 2024). Sampling at Lake Bourget was conducted over a two-day sampling campaign (i.e. 4^th^ and 5^th^ of September 2023), whereas all samples from Lake Paladru were collected in a single day (i.e. 28^th^ of September 2023).

Integrated and point sampling strategies (i.e. IS and PS respectively) were employed for eDNA collections in each lake (Figure 1). Integrated sampling consisted of collecting large water volumes (i.e. mean = 24 L, range = 8.5 – 40 L) along continuous littoral transects (i.e. maximum 10 m from the lake shore if navigation is possible) encompassing the entire lake shoreline (Civade et al., 2016; Hervé et al., 2022). A total of six littoral transects were sampled for Lake Bourget while two littoral transects were sufficient to cover the entire circumference of Lake Paladru (Figure 1). Integrated samples were collected by boat at a speed of *ca*. 5.5 km h^-1^ while continuously collecting sub-surface water using a diaphragm pump (Argaly, Sainte-Hélène-du-Lac, France) and filtering it through high capacity 0.45 µm Waterra capsules (Waterra, Mississauga, ON, Canada) for *ca*. 30 min. After filtrating, excess water was removed from the filter capsules and 40 mL (or 30 mL for one sample) of Longmire buffer was added to preserve eDNA samples which were stored at -80 °C upon return to the laboratory. Due to the greater depth of Lake Bourget, an additional deep-water sample was collected from the deeper northern basin (Figure 1). This involved pooling seven 5 L samples collected 5-10 m above the bottom substrate using a Niskin bottle, pooling of samples and filtering 29 L of water using a Waterra capsule. For PS strategies, three smaller volume (i.e. 2 L) samples were collected per littoral transect (i.e. at the start, middle and end of each transect) and for the deep-water sample (Figure 1). Point samples were collected using clean 2 L sampling bottles, preserved cooled and transported to the CARRTEL (INRAE – USMB) laboratory (Thonon-les-Bains) before filtering subsamples (i.e. mean = 1.7 L, range = 1.0 – 2.0 L) on the same day using 0.45 µm Sterivex filters (Merck KGaA, Darmstadt, Germany) and a vacuum pump (Vautier, 2024). Excess water was removed from the filters before adding 2 mL of Longmire buffer and storing samples at -80 °C until DNA extractions. A negative field control (NFC) was included for each sampling day. This control consisted of 2 L of MilliQ water in a clean bottle identical to those used for collecting point samples. The NFC was opened and closed on the boat to mimic field conditions and then filtered alongside the point samples.

### Molecular analyses

#### eDNA extractions

DNA was extracted from the filters using the NucleoMag DNA/RNA Water Kit (Macherey-Nagel, Düren, Germany) and the MagnetaPure 32 Nucleic Acid Purification System (Dutscher, Bernolsheim, France), with minor modifications to the manufacturer’s protocol. In an effort to enhance DNA recovery, a precipitation step was performed prior to extraction. The volume of preservation buffer used during the precipitation step varied by sampling type and consisted of the total volume of the recovered preservation buffer (from 1.7 mL to 2 mL) (Sterivex) or a 25 mL subsample (Waterra). Preservation solutions were extracted from the filter capsules and incubated overnight at 4 °C with 0.8 volumes of 100 % isopropanol, 0.2 volumes of 5 M NaCl, and glycogen (≥ 4.4 µg mL^-1^ final concentration). A 10 min centrifugation at 6500 g at room temperature was used to pellet the DNA and the supernatant was removed (Edmunds & Burrows, 2020). DNA pellets were resuspended in 550 µL of buffer MWA1 and subsequent extractions followed the protocol of Vautier et al. (2024). Total DNA was eluted in 120 µL of nuclease-free water, quantified using a NanoDrop^TM^ One/One^c^ spectrophotometer (Thermo Fisher Scientific, Waltham, MA, USA), and stored at -20 °C until further analysis. To control for potential contamination, six extraction blanks (i.e. unutilised Longmire Buffer) were processed alongside eDNA samples.

#### eDNA quantifications

Digital droplet PCR (ddPCR) utilizing the Bio-Rad QX600 system (Bio-Rad, Hercules, CA, USA) was employed to quantify eDNA concentrations of five target species and the entire fish community. Target species have historically been recorded in both lakes and included common roach (*Rutilus rutilus*), European catfish (*Silurus glanis*), perch (*Perca fluviatilis*), pike (*Esox lucius*), and whitefish (*Coregonus lavaretus*) (Jenny et al., 2024; STE, 2024). Species-specific assays for perch have previously been developed and validated (Vautier et al., 2023) while for all other target species assays were designed and validated in this study using the same workflow with a few modifications (see Supporting Information 1 (SI-1) and Table 1). The cytochrome c oxidase subunit I (COI) gene was targeted for all species-specific assays with ddPCR reactions containing 1x Bio-Rad ddPCR Supermix for probes (no dUTP) (Bio-Rad), 900 nM of each primer (Eurofins Genomics, Ebersberg, Germany), 250 nM of the internal probe (Eurofins Genomics), 5U AflII restriction enzyme (New England Biolabs, Ipswich, MA, USA), 3 µL template DNA, and nuclease-free water (Eurobio Scientific, Courtaboeuf, France) for a total volume of 22 µL. The primers and probes used to target each species are given in Table 1 and ddPCR reactions were performed in duplex, simultaneously targeting two species with probes containing two different fluorescent dyes (i.e. FAM and HEX). Droplets were generated using 20 µL of the PCR reaction mix with the BioRad QX200 Droplet generator (Bio-Rad). Amplification was achieved on a T100 Thermal Cycler (Bio-Rad) using an initial denaturation of 10 min. at 95 °C; 40 cycles of 30 sec. at 94 °C and 60 sec. at 60 °C and 10 min. at 98 °C. Quantification of the total fish eDNA concentrations was done using the generalist MiFish-U primers (Miya et al., 2015) with reaction mixtures containing 1x Bio-Rad ddPCR EvaGreen Supermix (Bio-Rad), 250 nM each of the MiFish-U primers (Eurofins Genomics), 3 µL template DNA, 5U AflII restriction enzyme (New England Biolabs), and nuclease-free water (Eurobio Scientific, Courtaboeuf, France) for a total volume of 22 µL. Thermal cycling conditions consisted of an initial denaturation for 5 min. at 95 °C; 40 cycles of 30 sec. at 95 °C and 30 sec. at 65 °C; 5 min. at 4 °C followed by 5 min. at 95 °C. Data analyses were performed using QX Manager software (Bio-Rad), with thresholds determined based on positive (DNA extracts from tissue samples) and negative (no-template extraction and ddPCR) controls and manufacturer’s instructions (Table 1). eDNA concentrations (copies µl^-1^ of DNA extract, copies µg^-1^ of total DNA extracted and copies l^-1^ of water filtered) were calculated while correcting for proportional volumes of preservation buffers used for DNA extractions.

**Table 1.**
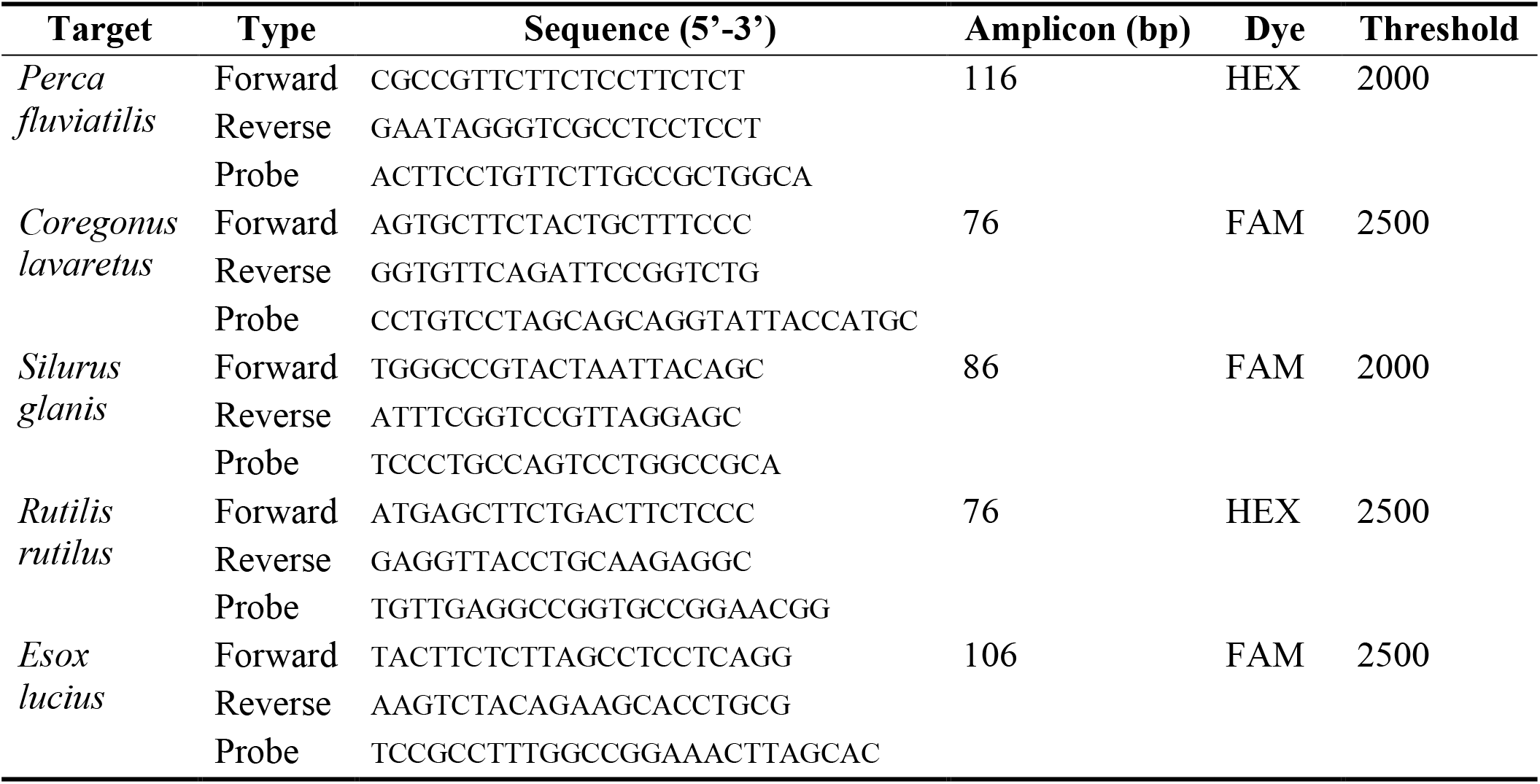
Details of the primers and probes used to quantify eDNA of the five target fish species.

### Statistical analyses

Statistical analyses and data visualisations were performed using R v4.2.3 (R Development Core Team, 2023) with the packages tidyverse (Wickham et al., 2019), glmmTMB (Brooks et al., 2017) and emmeans (Lenth, 2024). Linear and generalized linear mixed-effect models (i.e. LMM and GLMM respectively) were used for statistical analyses (Vautier et al., 2025). Additive models were fitted first and included the fixed effect terms of interest and random effects to account for the hierarchical data structure. Conditional Akaike Information Criterium (AICc) values (Akaike, 1974, 1978) were used to assess if interactions between fixed effects and the inclusion, when relevant, of a zero-inflation model improved the fit. Parameter significance (α = 0.05) was subsequently evaluated using ANOVA analyses (i.e. type III when interactions were present) and relevant pairwise comparisons through estimated marginal means and using Tukey p-value adjustments for multiple pairwise comparisons.

The effects of sampling strategies (i.e. PS vs. IS) on DNA recovery were assessed using the total DNA concentrations in the DNA extracts (ng µl^-1^), total fish DNA concentrations in the DNA extracts (copies µl^-1^) and fish DNA copies standardized by total DNA (copies µg^-1^ of total DNA extracted) as well as the fish DNA copies standardized by the volume of water filtered (copies l^-1^). For species-specific eDNA quantification only the latter measures (copies l^-1^) were used to assess the impact of sampling strategies on *in-situ* eDNA concentration estimates. Total DNA concentrations (ng µl^-1^) were transformed (i.e. Box-Cox transformation) to achieve normality and analysed using a LMM while fish DNA concentrations (copies µl^-1^, copies µg^-1^ and copies l^-1^) were analysed using a GLMM with a Negative-Binomial distribution. Most parsimonious models for total DNA and total fish DNA recovered included the full two-way interactions between sampling strategy and lake as fixed effects and transects nested within lakes as random effects. Effects of sampling strategies on species-specific eDNA quantifications were assessed excluding the results from Lake Paladru (i.e. low sample size) with the most parsimonious model including sampling strategy and species name as fixed effects, sampling transects as a random effect and a full additive zero-inflation model.

## RESULTS

One out of four NFCs tested positive for total fish DNA (i.e. 1 copy ul^-1^ of DNA extract), but this sample was negative for all target species. Consequently, the low DNA concentration observed in this negative control is likely attributable to off-target amplification of the MiFish-U primers which was confirmed through subsequent metabarcoding analyses (data not shown).

The IS strategy yielded significantly higher total DNA concentrations in Lake Bourget compared to all other lake-by-sampling strategies (Figure 2A). In contrast, in Lake Paladru the IS strategy significantly reduced the total fish DNA obtained compared to all other combinations (Figure 2B). When fish DNA copy numbers were standardized by total DNA and the volume of water filtered, the IS strategy showed a consistent lower recovery of fish DNA compared to the PS strategy, and these differences were significant in most cases (i.e. 3 out of 4 combinations) (Figure 2C and 2D).

**Figure 2.**
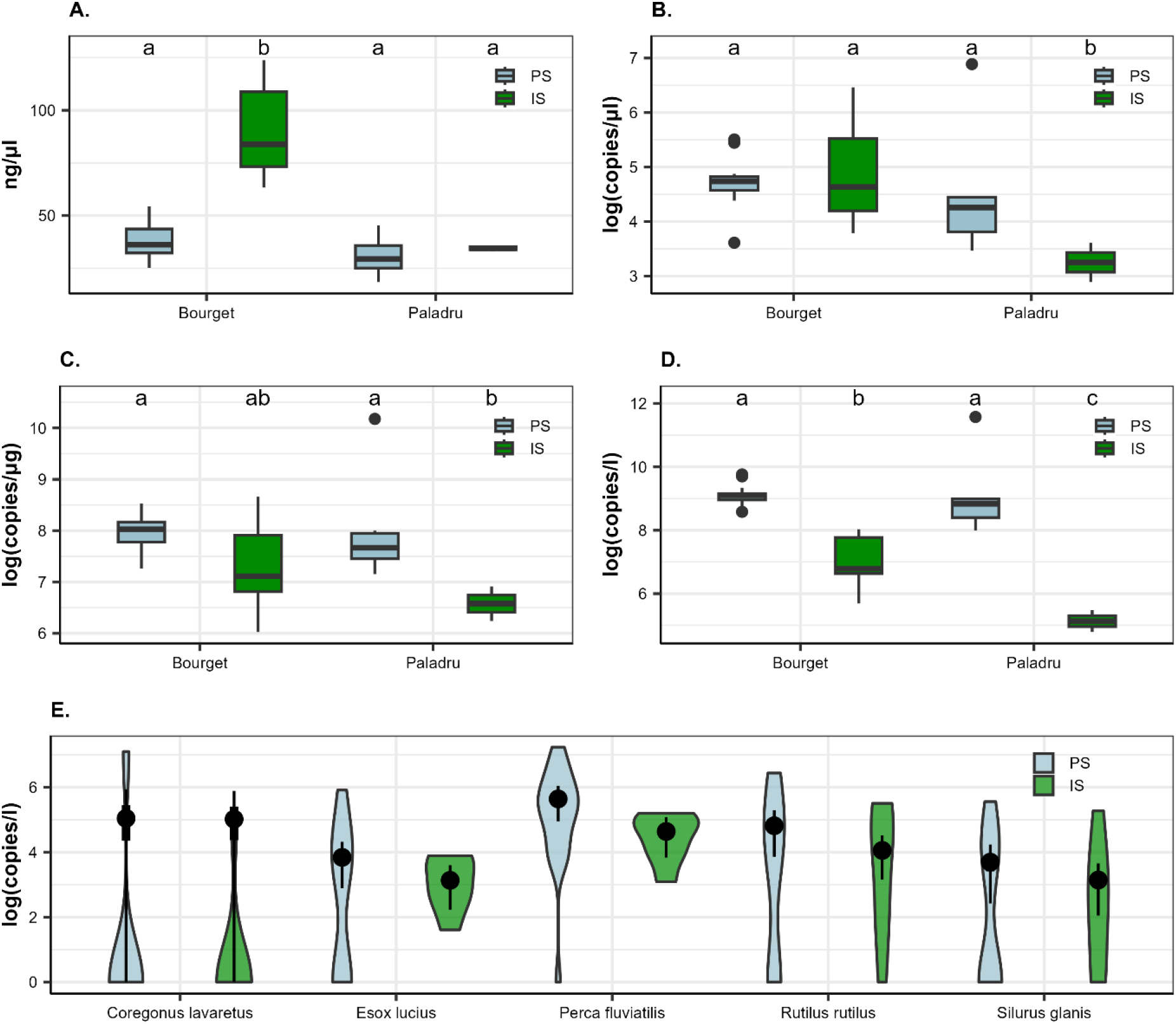
Effects of sampling strategies (i.e. point sampling (PS) or integrated sampling (IS)) for each lake on total environmental DNA (eDNA) concentrations in the DNA extracts (A), fish eDNA concentrations in the DNA extracts (B), fish eDNA copies standardized by total DNA extracted (C) or litre of filtered water (D), and eDNA concentrations per litre of filtered water for the five-target species in Lake Bourget (E). In panel E, solid points show the mean model estimates with the 50 % (thick lines) and 95 % (thin lines) confidence intervals.

Species-specific absolute eDNA quantifications were significantly affected by sampling strategy and species (SI-1). Parameter effects on the zero-inflation models also showed a significant effect of sampling strategy and species on the probability of zero detections (SI-1). Overall species-specific patterns were consistent with those observed for total fish DNA concentrations, with PS strategies yielding significantly higher eDNA concentrations, but simultaneously also increasing the probability for zero detections (i.e. violin plots are generally wider around zero copies l^-1^ for PS strategies) (Figure 2E). On average, the PS strategy resulted in species-specific eDNA concentrations that were 1.95 times higher compared to the IS strategy (range: 1.03 – 2.75). While rank abundances based on eDNA concentrations were largely similar between the two strategies, the dominant taxa differed between strategies, with perch eDNA concentrations being highest in the PS strategy, whereas whitefish eDNA concentrations were higher in the IS strategy (Figure 3).

**Figure 3.**
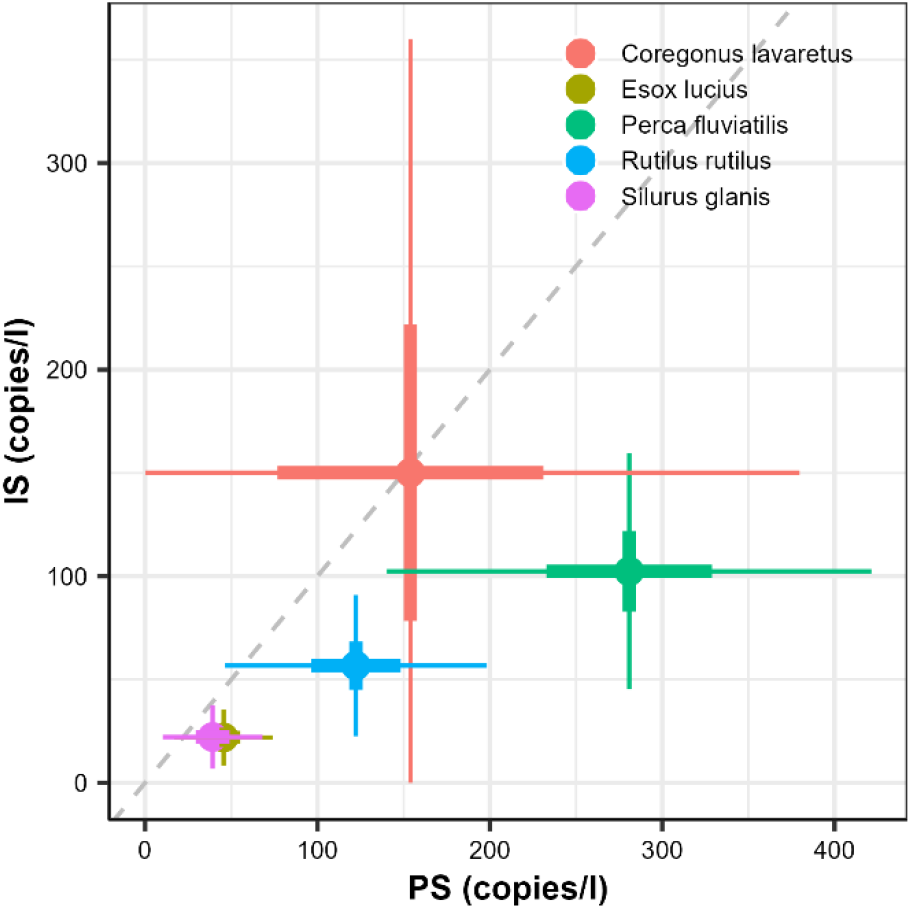
Absolute environmental DNA concentrations for the five target species in Lake Bourget observed with the point sampling (PS, n = 21) (x-axis) and integrated sampling (IS, n = 7) (y-axis) strategies. The grey dashed line showed the ideal 1 to 1 correlation.

## DISCUSSION

By implementing two alternative eDNA sampling strategies and performing absolute quantification of total fish eDNA and species-specific eDNA concentrations of five fish species, we assessed the performance of each sampling strategy to capture quantitative information. We found that collecting larger water volumes along littoral transects (i.e. integrated sampling) increased total DNA recovery but led to a reduced recovery of fish eDNA and lower *in-situ* fish eDNA concentrations compared to a point sampling strategy. The prevalence of zero detections was, however, higher for a point sampling strategy. These finding are partly congruent with previous results, which reported that filtering larger water volumes reduces DNA recovery per volume of water filtered and decreases detection probabilities for certain amphibian taxa (Peixoto et al., 2021).

Several, not mutually exclusive, explanations may account for these observations and will need to be evaluated in the future to understand the mechanisms at play. Firstly, the spatial coverage of the point sampling strategy may be insufficient, potentially requiring a greater number of point samples to accurately estimate mean eDNA concentrations (Yates et al., 2023). Indeed, we found that the variance associated with eDNA concentration estimates obtained from the point sampling strategy was higher compared to integrated sampling strategies (Figure 3). However, increasing sampling effort would not change the observation that total fish eDNA recovery, when standardized, is lower for the integrated sampling strategy. Secondly, the total water volume filtered may also affect DNA recovery with larger volumes leading to clogging and reducing the effectiveness of eDNA retention. However, when considering the ratio between the mean water volume filtered to filter area (i.e. 10 vs. 600 cm^2^ for Sterivex and Waterra filters respectively) values are higher for Sterivex compared to Waterra filters (i.e. 0.17 vs. 0.04) suggesting that clogging effects should be more pronounced for the point sampling strategy. Higher degradation rates of DNA for the integrated, relative to the point, sampling strategy may also explain the observed results. However, the on-site filtration and preservation used during the integrated sampling strategy should have provided for a better eDNA preservation. Point samples on the other hand were transported to the laboratory before filtration and preservation (*ca*. 4 hours between collection and filtration/preservation) allowing for more eDNA degradation. Differences between the sampling strategies may also simply be due to differences in the materials used in the filtering capsules. The point sampling strategy utilized polyvinylidene fluoride (PVDF) Sterivex filters, while the integrated sampling strategy relied on high capacity polyethersulfone (PES) Waterra filters. The former have been found to improve DNA recovery, although the effect may depend on the DNA extraction method used (Djurhuus et al., 2017; Min & Kim, 2024). Furthermore, magnetic bead-based DNA extractions have been shown to have a relative lower extraction efficiency compared to precipitation-based methods and DNA extraction efficiency may also vary depending on the state of eDNA collected (Kirtane et al., 2023; Kirtane & Deiner, 2024). A less effective DNA extraction for the Waterra filters may thus also explain the obtained results although the total eDNA concentrations observed in the integrated samples (i.e. maximum 124 ng/µL) suggest that a saturation, and subsequent lower DNA recovery, of the beads is unlikely. Filtering larger volumes may also increase the concentration of PCR inhibitors which may affect the ability to accurately quantify eDNA concentrations. However, magnetic bead-based DNA extraction have been reported to effectively remove inhibitors (Kirtane & Deiner, 2024) and inhibitors should not strongly influence DNA quantification through ddPCR (Capo et al., 2021; Doi et al., 2015).

We also found evidence that sampling strategies affect the rank abundance based on eDNA concentrations although this result is based solely on the results of one lake and five species. The dominance of perch eDNA in the point sampling strategy is congruent with standard catch per unit effort (CPUE) estimates, while a dominance of whitefish eDNA in the integrated sampling strategy is consistent with hydro-acoustic observations, which suggest a dominance of whitefish in Lake Bourget in terms of biomass (CPUE values and hydroacoustic data are presented in Jenny et al. (2024)). It is worth noting that whitefish eDNA concentration estimates from both sampling strategies do not vary as much as for other species. Of the species targeted, whitefish have a more pelagic lifestyle and occupy deeper waters while others are more strongly associated with the littoral habitat as adults. The differences between the rank abundance estimates between sampling strategies may thus be related to the fact that point sampling may capture a more local signal (i.e. higher perch abundance in the littoral zone) while the integrated sampling strategy may capture a more integrated signal (i.e. higher whitefish abundance in the lake).

Overall, we can conclude that the collection and filtering of spatially integrated large water volumes, possibly in combination with the DNA extraction methods used here, reduces fish eDNA recovery compared to a point sampling strategy involving the filtration of small volume of water. Disentangle the exact mechanisms leading to these differences in eDNA recovery and estimates of in-situ eDNA concentrations will require further examination using carefully designed study designs. Nonetheless our results show that a careful consideration of the implemented sampling strategy is needed depending on the objectives of the study (e.g. estimating local eDNA concentrations or maximizing species detections).

## ACKNOWLEDGMENTS

We wish to thank P. Perney and C. Galiegue for assisting with eDNA collections. C. Galiegue also contributed to the eDNA extractions and digital droplet PCR analyses.

## AUTHOR CONTRIBUTIONS

All co-authors contributed to the study design. MV performed the field and laboratory work with the assistance of JB. JB performed the statistical analyses and, MV and JB led the writing of the manuscript with significant contributions from all co-authors. All authors revised and approved the final manuscript for publication.

## DATA AVAILABILITY

The complete data and R script for the statistical analyses are available on the figshare data repository https://doi.org/10.6084/m9.figshare.30032209.v2 (Vautier et al., 2025).

