## Supplementary material for "Environmental DNA (eDNA) sampling strategies influence estimates of freshwater fish eDNA concentrations": SI-1

### Primer design and validation

Cytochrome oxidase I (COI) gene sequences for the five target species, as well as for 30 additional fish species present in French peri-alpine lakes (35 species in total; see Table S1), were retrieved from the Barcode of Life Database (BOLD; http://www.boldsystems.org/). All available sequences were included, regardless of geographic origin, to account for potential intraspecific genetic variation.

Primer and probe sequences targeting the COI gene were designed using Primer3 software (Untergasser et al., 2012). Design parameters for forward and reverse primers included a PCR product size range of 60–160 bp, primer melting temperatures between 50°C and 65°C (with a maximum difference of 3°C between primer pairs). Probe design parameters included a length of 18–27 nucleotides, melting temperatures between 60°C and 67°C, and a GC content of 30–80%.

To assess specificity, each primer set was tested *in silico* against the COI sequences of the 34 non-target fish species. Multiple sequence alignments were performed using MEGA7 (Kumar et al., 2016). Only primer/probe sets with mismatches to the non-target species were retained. Due to high sequence homology among certain species, the number of base-pair mismatches between each primer/probe and the non-target species was summed, and only those sets with the highest mismatch counts were selected (Table S1). The secondary structure of each amplicon was evaluated using the Mfold program (Zuker, 2003), and the specificity of each primer/probe set was further confirmed by performing primer BLAST analysis against the NCBI database to ensure exclusive targeting of the intended species.

**Table S1.** Total number of nucleotide differences between the primers/probe sets used in this study and the COI sequence of other fish species present in French peri-alpine lakes. In total, 35 fish species were tested, including the five target species (indicated by *).

| **Non-target species** | **Total number of base-pair mismatches with the primer/probe sets for the below target species** | | | | |
|---|---|---|---|---|---|
| | *Perca fluviatilis* | *Coregonus lavaretus* | *Silurus glanis* | *Esox lucius* | *Rutilus rutilus* |
| *Abramis brama* | 14 | 19 | 18 | 20 | 11 |
| *Alburnoides bipunctatus* | 20 | 18 | 16 | 21 | 10 |
| *Alburnus alburnus* | 13 | 17 | 14 | 21 | 14 |
| *Ameiurus melas* | 19 | 20 | 16 | 19 | 13 |
| *Barbatula barbatula* | 15 | 17 | 14 | 22 | 12 |
| *Barbus barbus* | 16 | 21 | 19 | 20 | 11 |
| *Blicca bjoerkna* | 17 | 18 | 16 | 19 | 10 |
| *Coregonus lavaretus** | 18 | 0 | 11 | 18 | 11 |
| *Cottus gobio* | 12 | 17 | 14 | 15 | 13 |
| *Cyprinus carpio* | 13 | 18 | 16 | 17 | 12 |
| *Esox lucius** | 15 | 14 | 13 | 0 | 12 |
| *Gasterosteus aculeatus* | 18 | 17 | 12 | 12 | 17 |
| *Gobio gobio* | 13 | 14 | 15 | 17 | 14 |
| *Gymnocephalus cernuus* | 14 | 16 | 15 | 23 | 15 |
| *Lampetra planeri* | 13 | 16 | 14 | 18 | 19 |
| *Lepomis gibbosus* | 18 | 19 | 18 | 17 | 15 |
| *Leuciscus leuciscus* | 16 | 14 | 10 | 19 | 15 |
| *Lota lota* | 15 | 17 | 18 | 18 | 11 |
| *Oncorhynchus mykiss* | 20 | 19 | 15 | 19 | 11 |
| *Perca fluviatilis** | 0 | 11 | 11 | 19 | 13 |
| *Phoxinus phoxinus* | 18 | 16 | 14 | 18 | 16 |
| *Pseudorasbora parva* | 15 | 16 | 14 | 21 | 11 |
| *Rhodeus amarus* | 18 | 13 | 15 | 16 | 12 |
| *Rutilus rutilus** | 14 | 14 | 13 | 17 | 0 |
| *Salaria fluviatilis* | 17 | 19 | 14 | 20 | 9 |
| *Salmo trutta* | 18 | 19 | 15 | 26 | 14 |
| *Salvelinus alpinus* | 18 | 14 | 14 | 17 | 15 |
| *Sander lucioperca* | 14 | 13 | 14 | 17 | 12 |
| *Scardinius erythrophthalmus* | 17 | 16 | 14 | 14 | 19 |
| *Silurus glanis** | 18 | 14 | 0 | 19 | 11 |
| *Squalius cephalus* | 16 | 13 | 14 | 14 | 10 |
| *Telestes souffia* | 16 | 18 | 18 | 19 | 9 |
| *Thymallus thymallus* | 15 | 16 | 15 | 19 | 9 |
| *Tinca tinca* | 11 | 15 | 12 | 13 | 12 |
| *Zingel asper* | 12 | 16 | 14 | 19 | 14 |

Experimental validation of primer/probe specificity was performed using droplet digital PCR (ddPCR) on DNA extracted from tissues of all 35 fish species (Table S1). For each assay, DNA pools were prepared containing 100 pg of DNA from all species except the target species. ddPCR reactions (22 µL total volume) consisted of 1× Bio-Rad ddPCR Supermix for probes (no dUTP), 900 nM of each primer (Eurofins Genomics, Ebersberg, Germany), 250 nM of the internal probe (Eurofins Genomics), 5 U AflII restriction enzyme (New England Biolabs, Ipswich, MA, USA), 4 µL pooled DNA, and nuclease-free water (Eurobio Scientific, Courtaboeuf, France). The ddPCR protocol followed the procedures described in the main manuscript. Primer/probe sets were considered specific only if no amplification was detected from non-target species in ddPCR assays.

### Statistical analyses

**Table S 2.** Summary table of the proportion of positive detections and the eDNA concentration in the DNA extracts (copies µl$^{-1}$) for the different sample types (i.e. negative extraction control (NEC), negative field control (NFC), negative template control (NTC), Sterivex filter capsules (Sterivex) and Waterra filter capsules (Waterra)) and target species.

| Sample type | Species | Gene region | Detections (pos/total) | Concentrations | | | |
|---|---|---|---|---|---|---|---|
| | | | | Mean | SD | Min. | Max. |
| NEC | All fish | 12S | 0/6 | 0.00 | 0.00 | 0 | 0 |
| NEC | *Coregonus lavaretus* | COI | 0/6 | 0.00 | 0.00 | 0 | 0 |
| NEC | *Esox lucius* | COI | 0/6 | 0.00 | 0.00 | 0 | 0 |
| NEC | *Perca fluviatilis* | COI | 0/6 | 0.00 | 0.00 | 0 | 0 |
| NEC | *Rutilus rutilus* | COI | 0/6 | 0.00 | 0.00 | 0 | 0 |
| NEC | *Silurus glanis* | COI | 0/6 | 0.00 | 0.00 | 0 | 0 |
| NFC | All fish | 12S | 1/4 | 0.25 | 0.50 | 0 | 1 |
| NFC | *Coregonus lavaretus* | COI | 0/4 | 0.00 | 0.00 | 0 | 0 |
| NFC | *Esox lucius* | COI | 0/4 | 0.00 | 0.00 | 0 | 0 |
| NFC | *Perca fluviatilis* | COI | 0/4 | 0.00 | 0.00 | 0 | 0 |
| NFC | *Rutilus rutilus* | COI | 0/4 | 0.00 | 0.00 | 0 | 0 |
| NFC | *Silurus glanis* | COI | 0/4 | 0.00 | 0.00 | 0 | 0 |
| NTC | All fish | 12S | 0/2 | 0.00 | 0.00 | 0 | 0 |
| NTC | *Coregonus lavaretus* | COI | 0/2 | 0.00 | 0.00 | 0 | 0 |
| NTC | *Esox lucius* | COI | 0/2 | 0.00 | 0.00 | 0 | 0 |
| NTC | *Perca fluviatilis* | COI | 0/2 | 0.00 | 0.00 | 0 | 0 |
| NTC | *Rutilus rutilus* | COI | 0/2 | 0.00 | 0.00 | 0 | 0 |
| NTC | *Silurus glanis* | COI | 0/2 | 0.00 | 0.00 | 0 | 0 |

| | | | | | | | |
|---|---|---|---|---|---|---|---|
| Sterivex | All fish | 12S | 27/27 | 138.00 | 174.57 | 31 | 979 |
| Sterivex | *Coregonus lavaretus* | COI | 5/27 | 2.44 | 5.77 | 0 | 19 |
| Sterivex | *Esox lucius* | COI | 13/27 | 0.85 | 1.29 | 0 | 5 |
| Sterivex | *Perca fluviatilis* | COI | 26/27 | 3.74 | 3.69 | 0 | 16 |
| Sterivex | *Rutilus rutilus* | COI | 19/27 | 2.26 | 3.16 | 0 | 13 |
| Sterivex | *Silurus glanis* | COI | 11/27 | 0.67 | 1.14 | 0 | 5 |
| Waterra | All fish | 12S | 9/9 | 170.67 | 211.92 | 17 | 640 |
| Waterra | *Coregonus lavaretus* | COI | 2/9 | 3.11 | 8.61 | 0 | 26 |
| Waterra | *Esox lucius* | COI | 7/9 | 2.33 | 2.40 | 0 | 8 |
| Waterra | *Perca fluviatilis* | COI | 9/9 | 10.00 | 8.14 | 2 | 29 |
| Waterra | *Rutilus rutilus* | COI | 8/9 | 7.56 | 7.09 | 0 | 18 |
| Waterra | *Silurus glanis* | COI | 5/9 | 2.67 | 4.24 | 0 | 11 |

### Total DNA concentrations in the DNA extracts (ng µl$^{-1}$)

**Table S3.** Anova summary table of the most parsimonious model.

| Term | $\chi^2$ | DF | P |
|---|---|---|---|
| (intercept) | 1.7 | 1 | 0.188 |
| Lake | 4.99 | 1 | ***0.025*** |
| Strategy | 56.06 | 1 | ***<0.0001*** |
| Lake:Strategy | 5.60 | 1 | ***0.018*** |

**Table S4.** Summary table of the pairwise comparisons between lake and sampling strategies (i.e. point sampling (PS) or integrated sampling (IS))

| Contrast | Estimate | SE | DF | P |
|---|---|---|---|---|
| Bourget PS - Paladru PS | 0.730 | 0.327 | 29 | 0.138 |
| Bourget PS - Bourget IS | -1.717 | 0.229 | 29 | ***<0.0001*** |
| Bourget PS - Paladru IS | 0.165 | 0.446 | 29 | 0.982 |
| Paladru PS - Bourget IS | -2.447 | 0.365 | 29 | ***<0.0001*** |
| Paladru PS - Paladru IS | -0.565 | 0.429 | 29 | 0.559 |
| Bourget IS - Paladru IS | 1.881 | 0.474 | 29 | ***0.002*** |

### Fish DNA concentrations in the DNA extracts (copies µl$^{-1}$)

**Table S5.** Anova summary table of the most parsimonious model.

| Term | $\chi^2$ | DF | P |
|---|---|---|---|
| (intercept) | 530.59 | 1 | ***<0.0001*** |
| Lake | 0.47 | 1 | 0.494 |
| Strategy | 2.36 | 1 | 0.124 |
| Lake:Strategy | 13.33 | 1 | ***<0.0001*** |

**Table S6.** Summary table of the pairwise comparisons between lake and sampling strategies (i.e. point sampling (PS) or integrated sampling (IS))

| Contrast | Estimate | SE | DF | P |
|---|---|---|---|---|
| Bourget PS - Paladru PS | -0.310 | 0.452 | Inf | 0.903 |
| Bourget PS - Bourget IS | -0.445 | 0.290 | Inf | 0.415 |
| Bourget PS - Paladru IS | 1.528 | 0.584 | Inf | ***0.044*** |
| Paladru PS - Bourget IS | -0.136 | 0.490 | Inf | 0.993 |
| Paladru PS - Paladru IS | 1.837 | 0.545 | Inf | ***0.004*** |
| Bourget IS - Paladru IS | 1.973 | 0.622 | Inf | ***0.008*** |

### Fish DNA concentrations standardised by total DNA (copies $ng^{-1}$)

**Table S7.** Anova summary table of the most parsimonious model.

| Term | $\chi^2$ | DF | P |
|---|---|---|---|
| (intercept) | 2637.54 | 1 | ***<0.0001*** |
| Lake | 2.19 | 1 | 0.139 |
| Strategy | 2.31 | 1 | 0.129 |
| Lake:Strategy | 7.15 | 1 | ***0.007*** |

**Table S8.** Summary table of the pairwise comparisons between lake and sampling strategies (i.e. point sampling (PS) or integrated sampling (IS))

| Contrast | Estimate | SE | DF | P |
|---|---|---|---|---|
| Bourget PS - Paladru PS | -0.547 | 0.369 | Inf | 0.449 |
| Bourget PS - Bourget IS | -0.417 | 0.275 | Inf | 0.426 |
| Bourget PS - Paladru IS | 1.445 | 0.480 | Inf | ***0.014*** |
| Paladru PS - Bourget IS | -0.963 | 0.400 | Inf | 0.075 |
| Paladru PS - Paladru IS | 1.991 | 0.508 | Inf | ***<0.001*** |
| Bourget IS - Paladru IS | 1.028 | 0.517 | Inf | ***0.192*** |

### Fish DNA concentrations standardised by total volume (copies $l^{-1}$)

**Table S9.** Anova summary table of the most parsimonious model.

| Term | $\chi^2$ | DF | P |
|---|---|---|---|
| (intercept) | 2430.90 | 1 | ***<0.0001*** |
| Lake | 1.98 | 1 | 0.160 |
| Strategy | 50.04 | 1 | ***<0.0001*** |
| Lake:Strategy | 21.03 | 1 | ***<0.0001*** |

**Table S10.** Summary table of the pairwise comparisons between lake and sampling strategies (i.e. point sampling (PS) or integrated sampling (IS))

| Contrast | Estimate | SE | DF | P |
|---|---|---|---|---|
| Bourget PS - Paladru PS | -0.587 | 0.417 | Inf | 0.495 |
| Bourget PS - Bourget IS | 1.871 | 0.265 | Inf | ***<0.0001*** |
| Bourget PS - Paladru IS | 3.976 | 0.523 | Inf | ***<0.0001*** |
| Paladru PS - Bourget IS | 2.459 | 0.452 | Inf | ***<0.0001*** |
| Paladru PS - Paladru IS | 4.564 | 0.517 | Inf | ***<0.0001*** |
| Bourget IS - Paladru IS | 2.105 | 0.558 | Inf | ***<0.001*** |

### Species-specific DNA concentrations standardised by total volume (copies $l^{-1}$)

**Table S11.** Anova summary table of the conditional part of the most parsimonious model.

| Term | $\chi^2$ | DF | P |
|---|---|---|---|
| (intercept) | 200.57 | 1 | ***<0.0001*** |
| Strategy | 27.78 | 1 | ***<0.0001*** |
| Species | 54.68 | 4 | ***<0.0001*** |

**Table S12.** Anova summary table of the zero-inflation part of the most parsimonious model.

| Term | $\chi^2$ | DF | P |
|---|---|---|---|
| (intercept) | 13.46 | 1 | ***<0.001*** |
| Strategy | 4.96 | 1 | ***0.026*** |
| Species | 27.42 | 4 | ***<0.0001*** |

**Table S13.** Summary table of the pairwise comparisons between sampling strategies (i.e. point sampling (PS) or integrated sampling (IS)) for the conditional model.

| Contrast | Estimate | SE | DF | P |
|---|---|---|---|---|
| PS - IS | 1.04 | 0.197 | Inf | ***<0.0001*** |

**Table S14.** Summary table of the pairwise comparisons between sampling strategies (i.e. point sampling (PS) or integrated sampling (IS)) for the zero-inflation model.

| Contrast | Estimate | SE | DF | P |
|---|---|---|---|---|
| PS - IS | 1.23 | 0.554 | Inf | ***0.026*** |
